# Motor cortical plasticity in response to skill acquisition in adult monkeys

**DOI:** 10.1101/2020.02.27.967562

**Authors:** Ankur Gupta, Abdulraheem Nashef, Sharon Israely, Michal Segal, Ran Harel, Yifat Prut

## Abstract

Cortical maps often undergo plastic changes during learning or in response to injury. In sensory areas, these changes are thought to be triggered by alterations in the pattern of converging inputs and a functional reassignment of the deprived cortical region. In the motor cortex, training on a task that engages distal effectors was shown to increase their cortical representation (as measured by response to intracortical microstimulation). However, this expansion could be a specific outcome of using a demanding dexterous task. We addressed this question by measuring the long-term changes in cortical maps of monkeys that were sequentially trained on two different tasks involving either proximal or distal joints. We found that motor cortical remodeling in adult monkeys was symmetric such that both distal and proximal movements can comparably alter motor maps in a fully reversible manner according to task demands. Further, we found that the change in mapping often included a switch between remote joints (e.g., a finger site switched to a shoulder site) and reflected a usage-consistent reorganization of the map rather than the local expansion of one representation into nearby sites. Finally, although cortical maps were considerably affected by the performed task, motor cortical neurons throughout the motor cortex were equally likely to fire in a task-related manner independent of the task and/or the recording site. These results may imply that in the motor system, enhanced motor efficiency is achieved through a dynamical allocation of larger cortical areas and not by specific recruitment of task-relevant cells.

## Introduction

Cortical plasticity is the ability of the cortex to reorganize itself in response to injury [1], changes in inputs [2–5] or during learning [6–9]. Although plasticity is often associated with the young brain, it has also been observed in adult animals and humans [10]. Studies of cortical plasticity in sensory areas in adults have shown that in response to partial sensory deprivation, cortical plasticity takes place in the form of changes in cortical maps [5]. It was initially assumed that these altered maps reflected changes in the representation of the periphery due to the modified pattern of convergence across afferent inputs [11, 12]. It appears, however, that the motor cortex is also capable of reorganization under specific circumstances. For example, training on a task that engaged finger movements induced the expansion of cortical fields related to distal muscles at the expense of proximal fields [13]. This result is consistent with recent findings in human subjects [14] showing that the organization of sensorimotor maps of the hands in humans reflects the daily use of fingers and not individual muscles. This use-dependent expansion of motor cortical maps was claimed to be a reflection of cortical recruitment of a larger number of synergies or the expanded size of individual synergies to allow for a more flexible and accurate implementation of the motor command required for controlling the working joints [15]. This hypothesis leads to several predictions: (1) the expansion in mapping of a task related joint should manifest in the takeover of nearby cortical sites which are likely to represent nearby joints; (2) the expansion process is non-symmetric; i.e., greater expansion is expected when acquiring distal as opposed to proximal skills, since rich sets of synergies are more critical for distal but not proximal motor skills; and (3) task-related activity should follow the plastic changes in the cortical map to take advantage of the modified control over motor output. This means that cell activity in the joint-relevant (expanded) sites should be more task-related than those in joint-irrelevant (contracted) sites.

To test these predictions, we examined the changes in motor cortical maps occurring after training on tasks that engaged either distal (fingers and wrist) or proximal (elbow and shoulder) joints and the neural correlates of the changes in the motor maps. The experimental paradigm consisted of sequentially training 3 monkeys on two types of target acquisition tasks: an isometric wrist task (WRIST) and a shoulder-elbow reaching task (REACH). We found that training monkeys on the WRIST task led to an expanded representation of distal joints in motor cortex, whereas monkeys trained on the REACH task had increased mapping of proximal joint representations. After retraining, single cortical sites expressed a change in output effects (from proximal to distal or the other way around) that was consistent with the different task requirements but often spanned more than a single joint. The extent of motor cortical plasticity (in terms of number of sites that changed their output effect and the joints spanned by changes in single sites) was insensitive to the order of the trained tasks. Finally, we found that the substantial changes in motor cortical mapping were not reflected in the magnitude of task-related activity of the motor cortical cells. These results suggest that the M1 map is modifiable in response to motor learning in adult monkeys. The observed plasticity was not limited to distal joints and can take place under different behavioral protocols. The changes in mapping were tightly linked to the performed task, and often span several joints, indicating a reorganization rather than expansion of the motor maps. Finally, the relations between the available motor maps and single cell firing patterns may imply that in the motor system, enhanced motor efficiency is achieved through recruiting more extensive cortical areas for output control and not by relying on specific activation of task-relevant cells.

## Results

### Changes in motor cortical mapping after training on different tasks

Monkeys (n=3) were sequentially trained on an isometric wrist task (WRIST, Fig. 1A) and shoulder-elbow reaching task (REACH, Fig. 1B). Both tasks required the monkey to wait for a go signal and then either to generated a directional torque to acquire the previously cued target or to make a reaching movement toward a peripheral target. The order and duration of training and recordings periods varied across monkeys (Fig. 1C-E). We mapped 693 cortical sites after WRIST training and 550 sites after REACH training (see Table 1 for monkey-specific numbers). We only considered sites that produced responses in the contralateral upper-limb (fingers to upper back). Figure 2 presents the motor cortical maps obtained for the three monkeys after training on the WRIST (upper row) and REACH (lower row) tasks. A number was assigned to each joint (1-fingers, 2-wrist, 3-elbows and 4-shoulder, 5-back). In cases where a single site was mapped more than once, we computed the median of the joint representation(using the scores for the joint representation). The joint-association of each site is color-coded from red (finger sites) to blue (shoulder sites). There was a general tendency for a prevalence of distal sites after WRIST training and proximal sites after REACH training. Comparing the distribution of single joint representations on the WRIST and REACH tasks revealed consistent changes between the maps obtained after each training period (Fig. 3). Here we only considered evoked responses of fingers, wrist, elbow and shoulder which constituted the vast majority of observed responses. The responses evoked by motor cortical stimulation obtained after WRIST training (Fig. 3A) showed an enhanced representation of the distal joints (finger and wrist, 62% of the total sites) whereas after REACH training (of the same monkeys) the representation of proximal sites increased (elbow and shoulder, 63%). This was true when considering sites obtained throughout the motor cortex as well as when focusing on the primary motor cortex (Fig. 3B, M1, 72% and 56% respectively). The changes in motor cortical mapping were significant when considering sites across the motor cortex (χ^2^ = 26.1, df=3, p < 9.2e-6) and when only considering sites in M1 (χ^2^= 25.7, df=3, p < 1.1e-5). We also examined the motor cortical maps of two additional monkeys trained on a single task alone (Fig. 3C-D). For the monkey trained on the WRIST task (orange bars in Fig. 3C-D) we found a larger finger and wrist representation (84 % in MC and 90% in M1) whereas in the monkey that was trained on the REACH task (blue bars in Fig. 3C-D), the proximal representation increased (71% in MC, 64% in M1). Comparing the joint representation between the two monkeys revealed significant differences for the motor cortex data (Fig. 3C, χ^2^= 78.0, df=3, p < 1.1e-16) and when considering M1 sites alone (Fig. 3Dχ^2^= 56.4, df=3, p < 3.4e-12). Table 1 shows the single monkey data.

**Figure 1:**
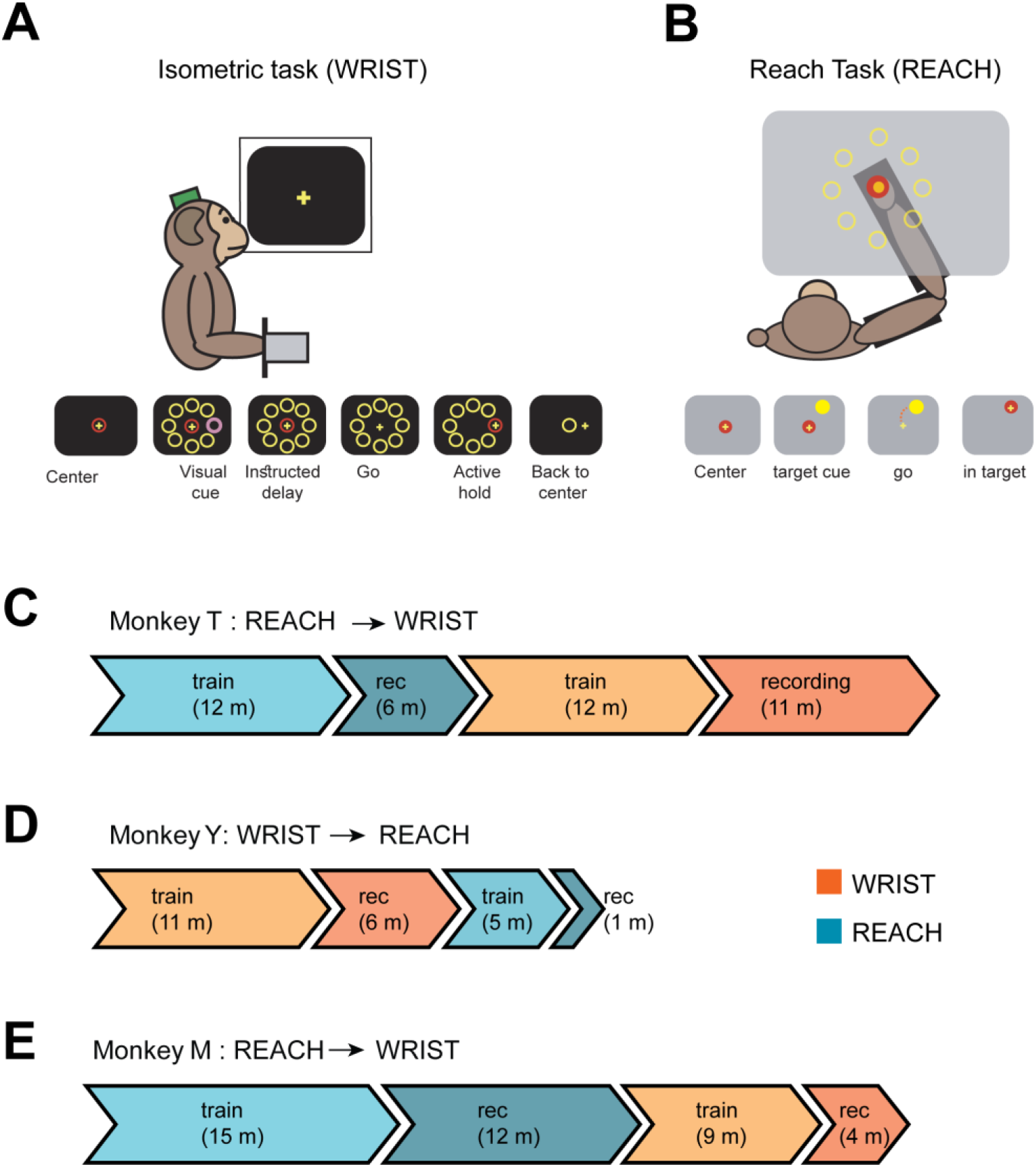
Behavioral paradigm and training time lines. **(A)** Schematic illustration of the isometric wrist task (WRIST). The monkey controlled an on screen cursor by generating a 2-dimensional wrist torque. The sequence of events composing a single trial is shown below. (**B)** Illustration of the setup configuration and sequence of events composing a single reach (REACH) trial. (**C)** Timeline of training and recording period (specified in months) for Monkey T. This monkey was first trained on the reach (REACH) task and then was retrained to perform the isometric wrist task (WRIST). Exact time durations for each training and recording period are indicated and the bar width is proportional to the time period of each phase. **(D)** Same as C but for monkey Y that was first trained on the wrist task and then on the reach task. **(E)** Same as C but for monkey M that was first trained on the reach task and then on the wrist task.

**Figure 2:**
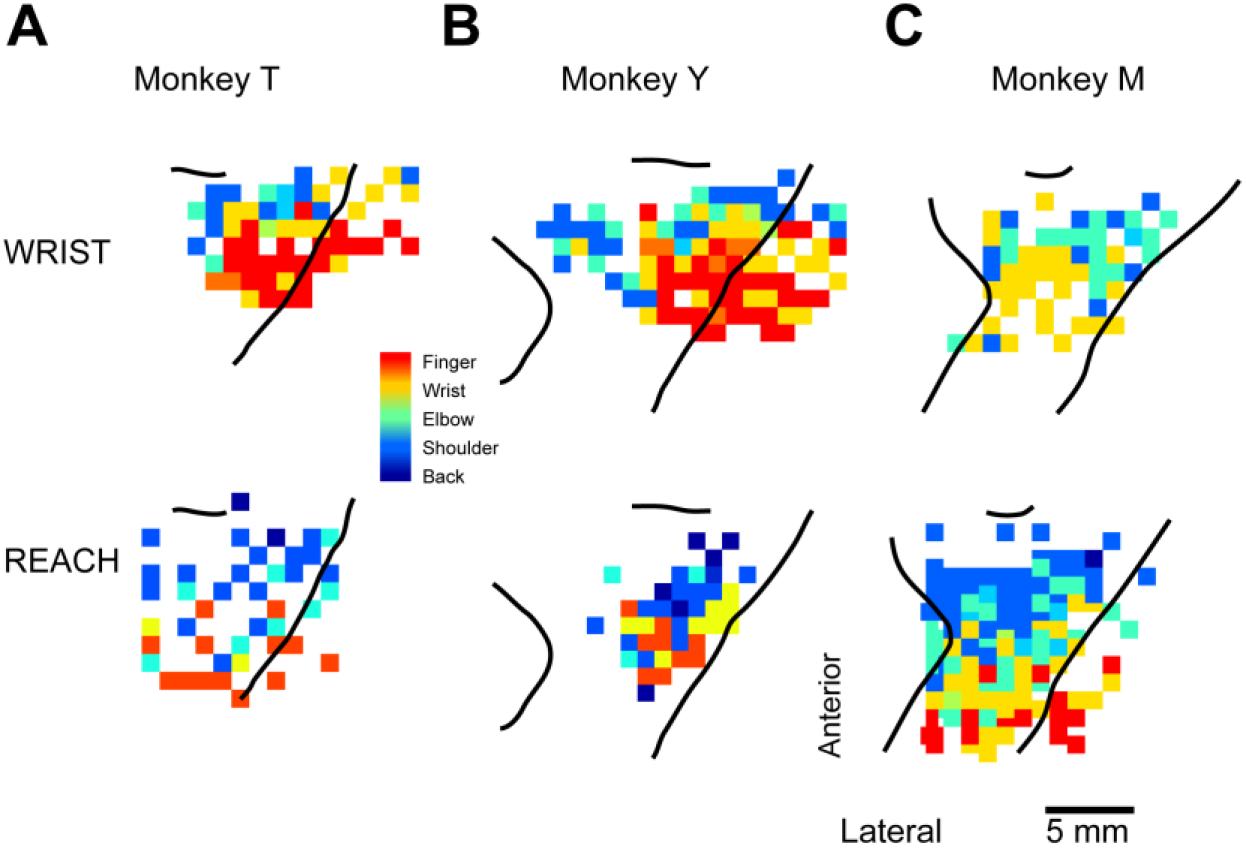
Representation of arm-related areas in motor cortex. Cortical maps obtained by inspecting joint movements evoked by intracortical microstimulation at each cortical site. The upper limb representation is comprised of digit (red), wrist (yellow), elbow (green), shoulder (light blue) and back (dark blue) movements. Top row presents the cortical map obtained after training on the wrist task for monkey T (A), monkey Y (B) and monkey M (C). Bottom row shows the cortical maps obtained for the same monkeys after training on the reach task.

**Figure 3:**
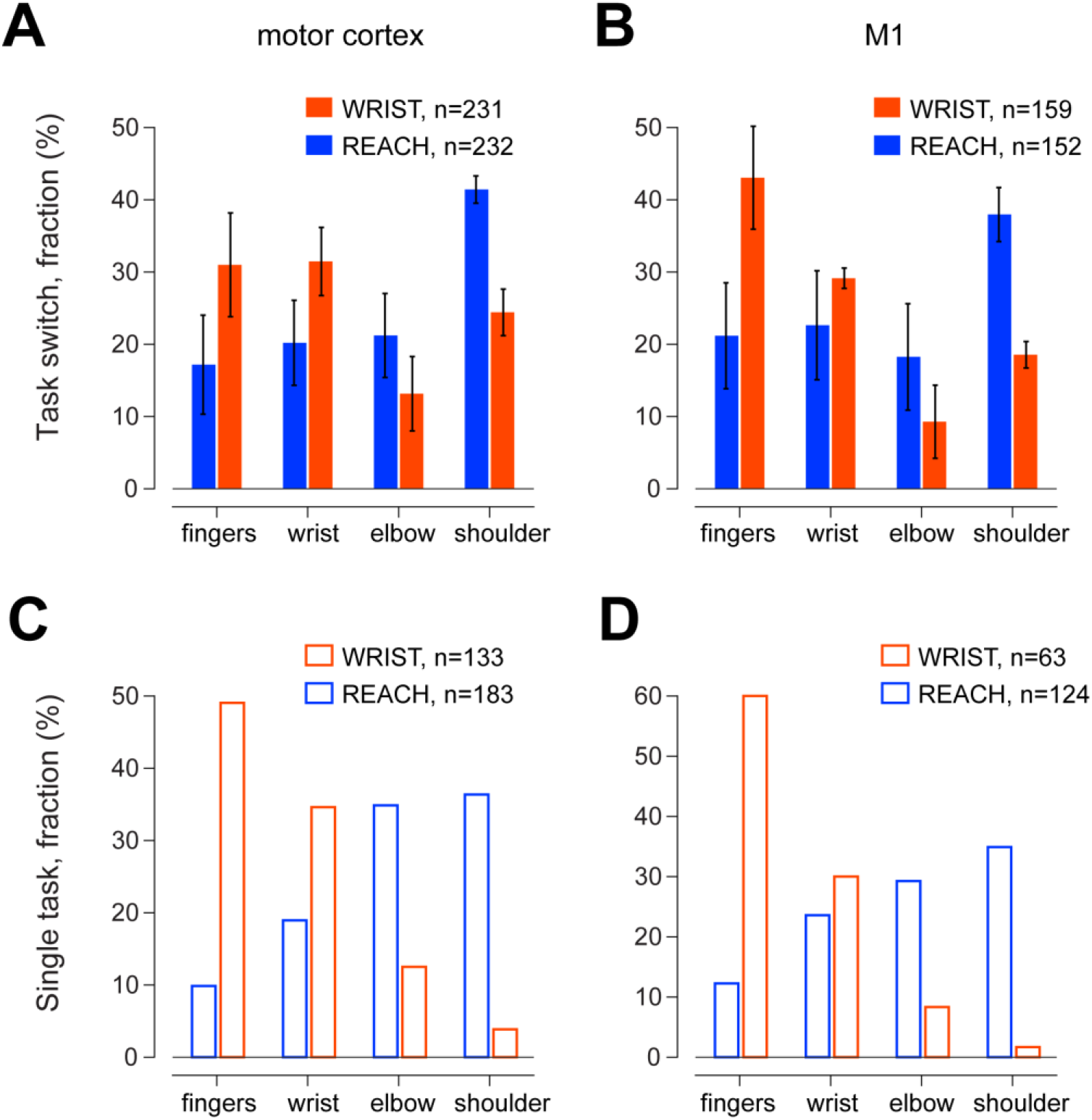
Task-specific joint representation in motor cortex. (A) Distribution of evoked responses obtained for cortical sites mapped in REACH (blue bars) and WRIST (orange) tasks. Data obtained for the motor cortex in the three monkeys tested on both tasks (sequentially). (B) Same monkeys as in A, but only considering sites in the primary motor cortex, M1. (**C**) Data obtained from the motor cortex of two monkeys which were trained on a single task. The orange bars (WRIST task) represent data obtained from one monkey (monkey V) and the blue bars represent data obtained from the other monkey (Monkey C). (**D**) Same data set as in C, but only considering M1 sites. Error bars represent SEM.

**Table 1:**
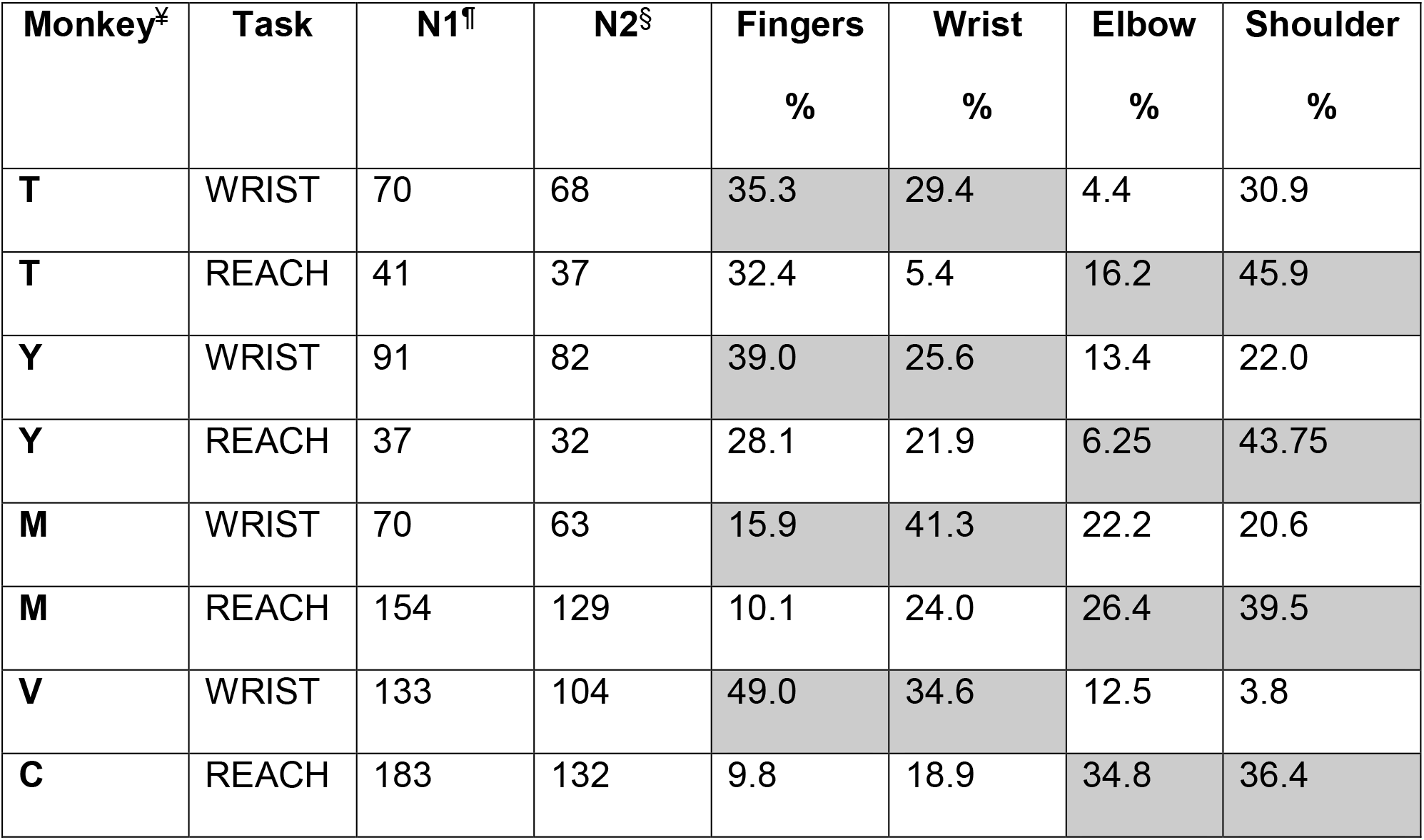
Number of mapped sites. ¥ - Monkey T, Y and M were recorded on ISO and KIN tasks. Monkeys V and C were recorded on a single task. ¶ - Total number of sites in motor cortex §-Total number of sites in motor cortex in which stimulation evoked response in the fingers, wrist, elbow or shoulder.

| Monkey<sup>¥</sup> | Task | N1<sup>¶</sup> | N2<sup>§</sup> | Fingers<br>% | Wrist<br>% | Elbow<br>% | Shoulder<br>% |
| --- | --- | --- | --- | --- | --- | --- | --- |
| **T** | WRIST | 70 | 68 | 35.3 | 29.4 | 4.4 | 30.9 |
| **T** | REACH | 41 | 37 | 32.4 | 5.4 | 16.2 | 45.9 |
| **Y** | WRIST | 91 | 82 | 39.0 | 25.6 | 13.4 | 22.0 |
| **Y** | REACH | 37 | 32 | 28.1 | 21.9 | 6.25 | 43.75 |
| **M** | WRIST | 70 | 63 | 15.9 | 41.3 | 22.2 | 20.6 |
| **M** | REACH | 154 | 129 | 10.1 | 24.0 | 26.4 | 39.5 |
| **V** | WRIST | 133 | 104 | 49.0 | 34.6 | 12.5 | 3.8 |
| **C** | REACH | 183 | 132 | 9.8 | 18.9 | 34.8 | 36.4 |

### Site-specific changes in mapping

After finding a training-induced and task-consistent change in mapping properties across the motor cortex, we tested the extent of change in mapping properties in single cortical sites. To this end, we focused on single sites (n=228) in which mapping information had been collected for both the REACH and WRIST tasks (Fig. 4A). We first tabulated the frequency of each possible change in single-site mapping obtained between the two tasks (Fig. 4B). The results showed that roughly one-third of the tested sites (28%) in all monkeys maintained their mapping properties (gray-colored diagonal on the matrix in Fig. 4B). For the remainder of the sites where mapping properties changed, we defined a *task consistent* change (green cells in Fig. 4B) to be a change in mapping which corresponded to the switch in tasks performed (i.e., a distal joint site becoming a proximal joint site in a monkey which was first trained on the WRIST and then the REACH task). Likewise, a *task-inconsistent* change (pink cells in Fig. 4B) was defined as a change in mapping in the opposite direction to the switch in the order of the tasks (i.e., a distal joint site becoming a proximal joint site in a monkey that switched from the REACH to the WRIST task). The results showed that 79% percent (out of all the labile sites) had a task-consistent change in mapping, whereas only 21% exhibited an inconsistent change in mapping (a ratio of 3.8 fold).

**Figure 4:**
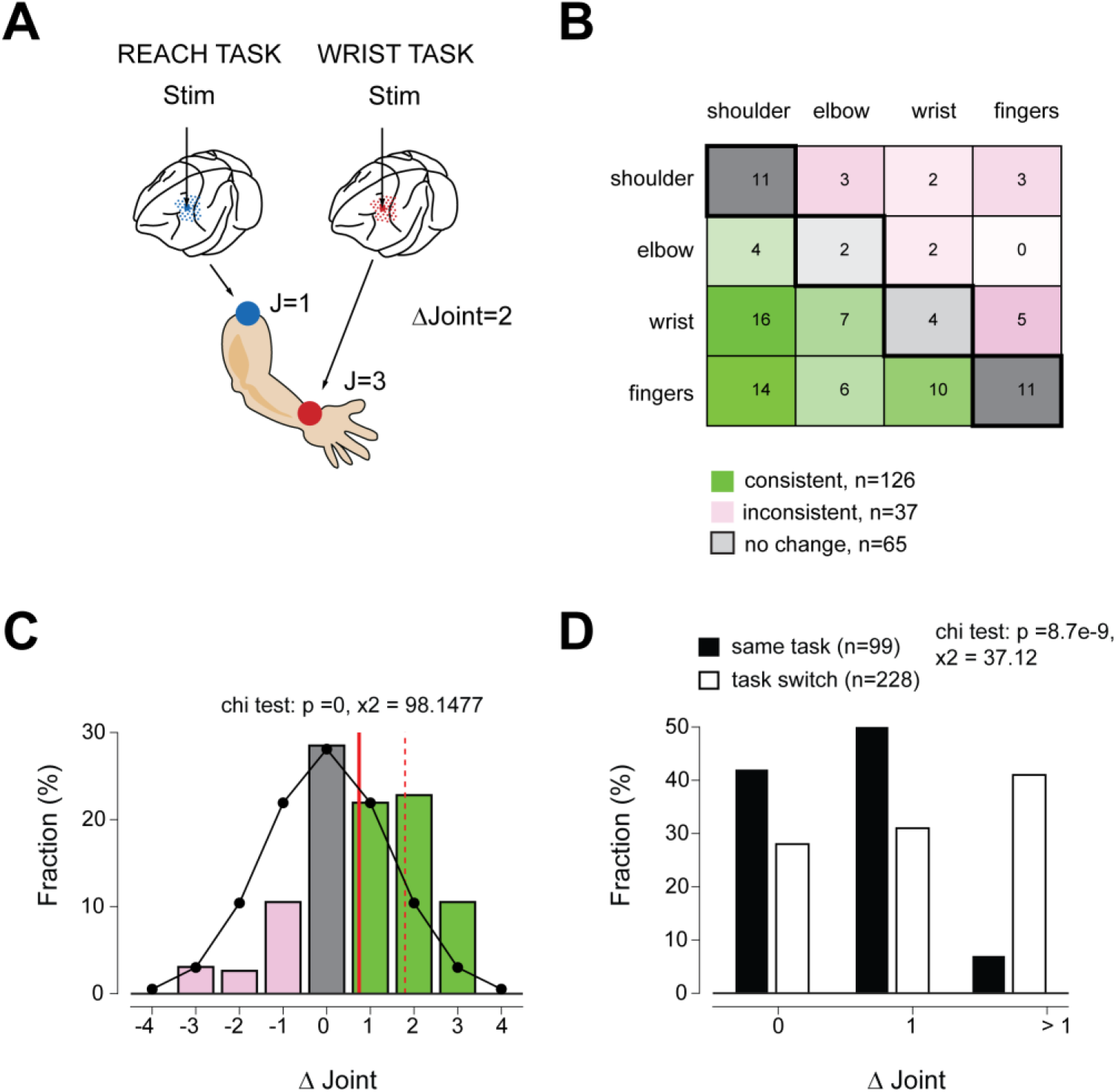
Task-dependent changes in mapping of single cortical sites. (A) Illustration presenting the way the joint-based metric was defined. We assigned a numerical value to each joint (shoulder=1, elbow=2, wrist=3, fingers=4). In the example, stimulation in a single cortical site produced a shoulder response on one task and a wrist response after switching to the second task. The inter-mapping distance was defined as ∆J=3-1=2. (B) For each possible combination of joint pairs, we computed the fraction of sites in which the change in mapping (before and after the task switch) corresponded to this pair. We further distinguished cases where this joint combination occurred in a consistent manner with the switch in task (green) and the fraction of sites in which the change in mapping was inconsistent with the switch in tasks (pink). Values along the diagonal (gray squares with thick outlines) represent cases in which the site had the same mapping properties after task switch. The intensity of each cell in the matrix corresponds to the fraction of sites found for the specific combination (specific percentages, computed as the average fraction across the three monkeys, are indicated in each cell). (C) Distribution of inter-site joint distances for all three monkeys combined (by summing the single monkey count data and computing fractions for the pooled dataset). Color of bars is according to the conventions in A. The vertical red line is the mean joint distance across the entire dataset and the dashed red line is the mean value for sites in which ∆J >0. Black line shows the expected distribution when assuming mean ∆J=0. (D) Distribution of absolute ∆J values obtained when testing the same site after task switch (white bars) or on the same task (black bars) was obtained when testing the same site multiple times while the monkeys continuously performed the same task.

Next, we calculated the distribution of inter-joint distances between the two maps (Fig. 4C). For this we defined a “joint-distance” metric by first assigning an index number (finger =1, wrist=2, elbow=3, shoulder=4) to each joint and then calculating the inter-task joint distance as ∆J_i_= (J_i1_-J_i2_) where J_i1_ is the index number for the *i-th* site as measured after the 1^st^ training period and J_i2_ is the index number for the same site but obtained after the 2^nd^training period. For instance, a site for which the same joint response was obtained on the two tasks corresponded to ∆J = 0, whereas a site for which stimulation evoked a shoulder response on one task and a wrist response on the second task yielded a distance of ∆J =3-1=2. The sign of the distance value (positive or negative) reflected a task-consistent or task-inconsistent change in mapping as was defined for Fig. 4B. The distribution of distances computed for the entire dataset (Fig. 4C) was found to be right-shifted towards positive values and was significantly different from the expected counts when assuming that the mean shift was zero (X^2^=98.1, df=6, p << 0.001). In addition, when considering only those sites in which joint mapping switched in a consistent manner (i.e., ∆J > 0) the mean distance was 1.8, with more than half of the sites having ∆J > 1 (76 vs. 50 sites). Finally, we compared the inter-joint distances obtained for single sites after the task switch to the distribution of distances obtained when mapping single sites multiple times while monkeys performed the same task (Fig. 4D) a value which should correspond to the “noise” in the mapping process (when the task is unchanged). There was a significant difference between the two distributions (X^2^ = 37.1, df =2, p < 9e-9) with a clear shift towards larger inter-joint distances when switching the task compared to the mapping during the same task. Taken together, these results suggest that training on a new task induces a task-consistent reorganization of the motor map rather than a simple expansion of the individual joint representation.

### Relationships between cortical maps and task-related properties of neurons

After documenting the changes in mapping that took place in response to and in correlation with motor skill learning, we further examined the cell activity correlates of the observed map remodeling. Our underlying hypothesis was that the increased number of task-relevant sites should be accompanied by a higher fraction of task-related neurons in these areas compared to task-irrelevant sites. The rationale behind this prediction was that this enhanced task-related activity restricted to joint-relevant cortical sites would better utilize the expanded representation for a more efficient and accurate recruitment of output effectors. To test this hypothesis, we analyzed the activity of single cortical neurons recorded from cortical sites with different mapping properties. The activity of each cell was tested for task-related modulation and directional tuning in the peri-movement (or torque onset) epoch. Figure 5 shows examples of cell pairs recorded simultaneously when monkeys performed the REACH task that relied on shoulder-elbow movements (Fig. 5A-B) and in a monkey performing the WRIST task which relied on wrist-finger movements (Fig. 5C-D). All 4 neurons were task-related and directionally tuned, irrespective of the output properties of the specific sites of the neurons and the performed task. For instance, the neurons in Fig. 5A and B were recorded simultaneously while the monkey performed the REACH task. Both neurons were task-related and tuned even through one was recorded from a shoulder site (Fig. 5A) and the second (Fig. 5B) was recorded from a finger site. Similarly, the two neurons that were recorded simultaneously while the monkey performed a WRIST task (Fig. 5C-D) were both task-related and tuned although they were recorded from very different sites. When quantifying this tendency across 5 monkeys for which we had both single unit activity and mapping information, we found that the fraction of task related and/or directionally tuned cells was constant irrespective of the performed task and/or the joint-identity of the recorded sites (Fig. 6A). The number of task related (or tuned) cells was not significantly different from the expected counts based on the assumption of a constant fraction of task-related neurons. This implies that in practice, the fraction of task-related (or tuned) cells remained constant across the entire hand-related motor area irrespective of the joints used in the specific tasks.

Next, we compared the firing and response properties of tuned cells recorded from sites with different output effects (proximal or distal joints) and during different tasks (REACH or WRIST). We found that the baseline firing rate of the cells (i.e., firing in the pre-cue period) and the peak firing rate of cells at the preferred direction were not sensitive to the mapping properties (Fig. 6B-C). Only the baseline level of cells recorded from proximal sites during the REACH task was higher than the level of cells recorded during the same task but from distal sites (Fig. 6B, p <0.02). Finally, we found that the average response profile of cells exhibited a similar pattern across the motor cortex (Fig. 6DE). Taken together, these results suggest a non-specific (i.e., site-independent) control of task-related activity and response patterns throughout the motor cortex.

**Figure 5.**
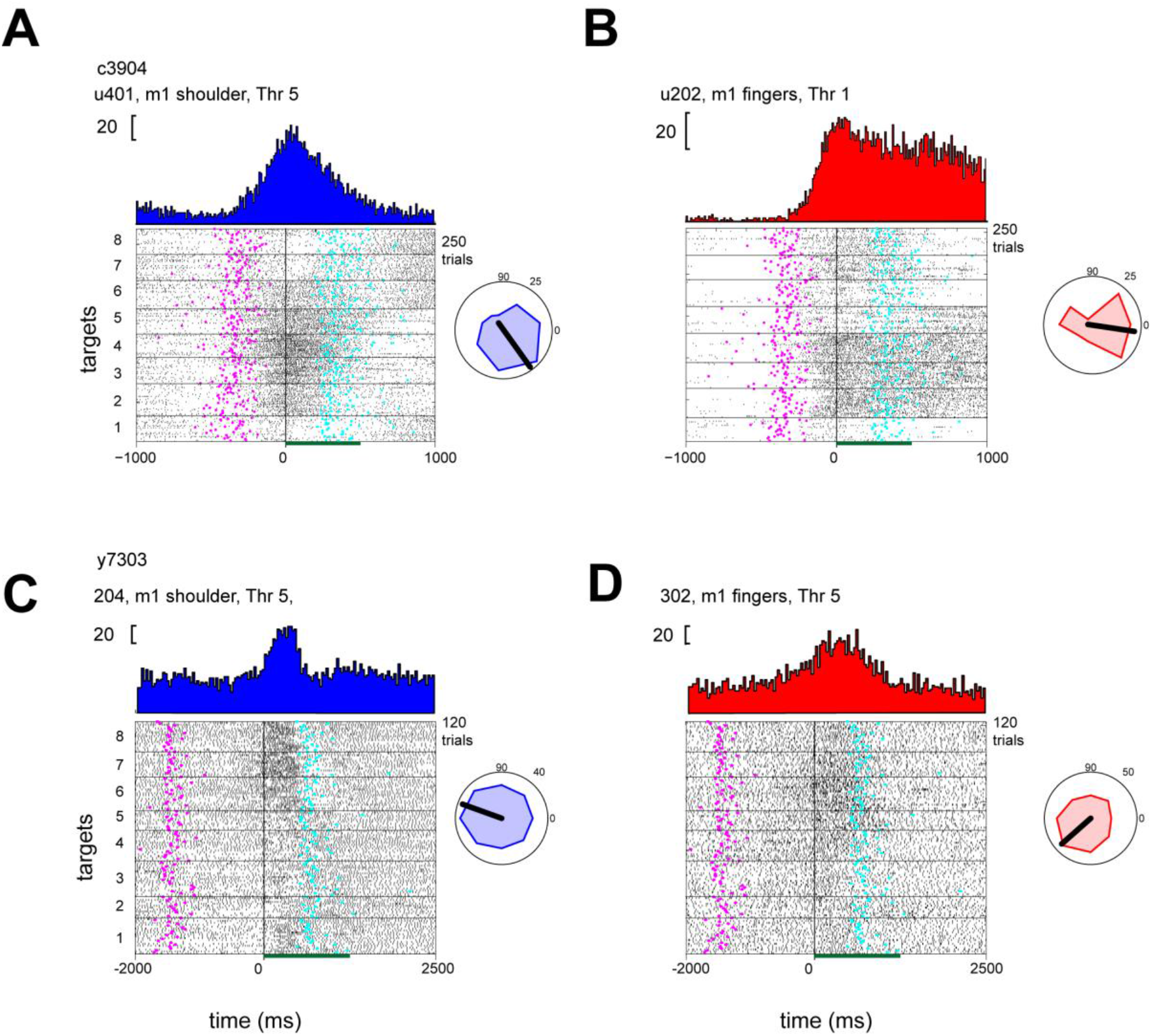
Site-dependent activity of simultaneously recorded cells. (A) Raster plot and PETH of a single unit recorded in a shoulder-related site of M1 during a REACH task. Trials are sorted by target number (1 to 8) and aligned on movement onset (t=0). PETH was computed using 25 ms bin size. Magenta and cyan symbols correspond to the time of cue-onset and target acquisition respectively on each trial. Threshold for joint activation (Thr) was 5 uAmp. (B) same as in A, but for a unit that was recorded at the same time from a different electrode which was located in a finger-related site of M1. Threshold for joint activation (Thr) was 1 uAmp. (C) Unit recorded during a WRIST task from a shoulder-related site in M1. (D) Unit recorded simultaneously as the unit in C but from a different electrode located in a finger-related site.

**Figure 6.**
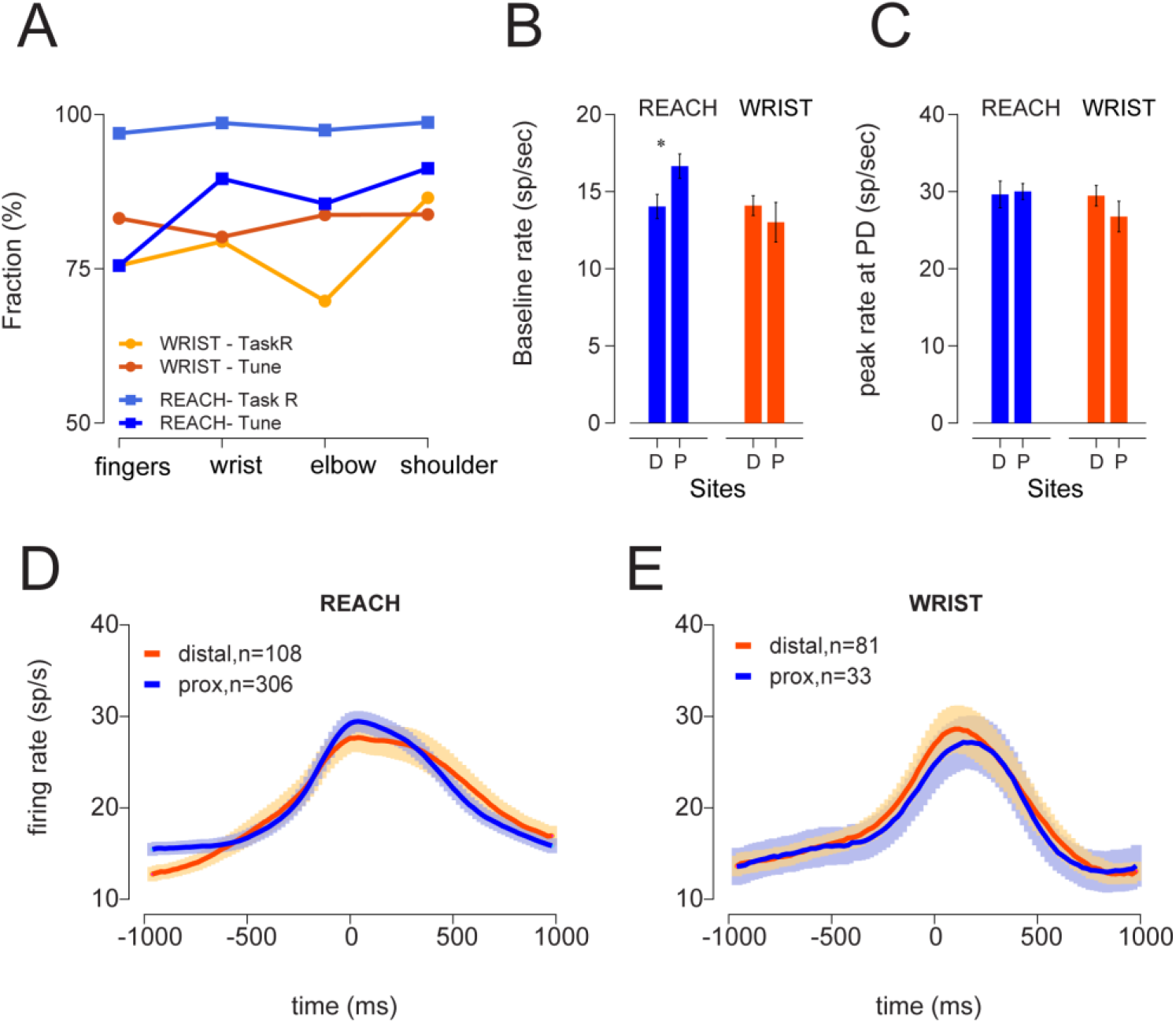
Quantification of response pattern as a function of site and task. (A) Fraction of task-related (TaskR) and tuned (Tune) cells found for each task (WRIST or REACH) at different sites (sorted by mapping properties). (B) Mean baseline firing rate computed for cells recorded in REACH (blue) or WRIST (orange) tasks from distal-related (D, fingers/wrist) or proximal-related (P, elbow/shoulder) sites. A significant difference in mean rate was found for REACH task (Kruskal Wallis task for medians, p <0.02). (C) Same as B but for peak rate at the preferred target for all cells. No significant differences were found between rates in either task. (D) Mean response pattern obtained for cells recorded on the reach (REACH) task from distal-related sites (orange) and proximal-related sites. Only tuned cells from M1 were considered here. (E) Same as in D but for cells recorded on the wrist (WRIST) task. Here we only considered data from monkey Y and T since the task parameters of monkey V were different.

## Discussion

Cortical plasticity is considered to be a fundamental brain process which often reflects changes in inputs in sensory systems or in use in motor systems [8, 10, 16]. The ability of the adult brain to change in response to normal or pathological perturbations attracts considerable attention because harnessing this capacity could contribute to the recovery from debilitating brain injuries [1, 17, 18]. Previous studies have shown that after training in finger movements, the finger related area in motor cortical sites expanded at the expense of proximal joint representations [13]. The current study extends this study by showing that in the adult brain the remodeling is symmetric in that both distal and proximal movements can comparably alter the representational maps in a reversible manner so that the modulation of maps under training in one task can be reversed following a training in a second motor task. Second, we found that the change in mapping reflected a usage-consistent reorganization of the map rather than the local expansion of one representation into nearby sites. Finally, the results showed that the substantial changes in mapping of the effectors were not accompanied by similar changes in the fraction of task-related and tuned cells across the motor cortex.

A critical first step in the formation of appropriate motor commands is the selection of appropriate motor primitives (i.e., muscle synergies) from the limited group of available synergies [15, 19–23]. Successful performance of skilled movements are then reinforced and the motor cortex undergoes network level changes [24–26]. Importantly, neuroplasticity has been shown to occur during reward-dependent skill learning and not when repeatedly performing a motor task [27, 28]. What are the functional implications of this increased representation in the motor domain? Initial studies focused on the expansion of distal joints after the training of naïve animal, with the underlying assumption that the change in the motor map reflected increased control over multi-joint motor tasks [13]. Here we found that motor plasticity was bidirectional, in that either the proximal or the distal joint maps expanded after appropriate training. It thus appears that the expansion of joint representations is not a mere reflection of task complexity, since switching from a distal task to a proximal test induced a similar reorganization of the maps even though the task complexity decreased. It is possible that in the intact motor cortex, while learning a new skill, the existing synergies are insufficient and lead to suboptimal movements. More efficient performance (in terms of the ultimate reward) can be achieved by recruiting a larger number of cortical sites which correspond to an expanded array of finely tuned muscle synergies [14, 29, 30]. In this scheme, task complexity alone does not dictate the expansion of the map; rather, the reward-based drive to optimize performance triggers map reorganization when acquiring either proximal or distal tasks, as a function of task requirements.

When switching the motor task, a significant fraction of the sites showed a change in joint mapping that was greater than 1 unit in the direction of the joints recruited for the final task. Previous studies have suggested that the plasticity of maps in the somatosensory area is based on the expansion of more active joints at the expense of adjacent under-active joints in a way consistent with the somatotopic representation [3, 4]. It is possible that the change in representation in the motor cortex reflects an underlying need for better control of the performed actions and not a modification of the local sensory inputs [31, 32].

The changes in motor cortical maps were not accompanied by corresponding changes in the fraction of task-related and/or tuned cells, indicating that throughout the motor cortex, a large fraction of neurons (recorded from sites that controlled task-related as well as task-unrelated joints) exhibited comparable task-related activity. This pattern is consistent with the “dense” coding scheme employed in motor and premotor areas [33, 34], where many cells exhibit task related activity. This is a very different operational principle than the sparse coding scheme often ascribed to sensory cortical areas [35, 36]. Our data revealed that response patterns and firing properties of single cells were similar across the motor cortex irrespective of the output site and the performed task. This result is consistent with earlier studies that have reported the absence of single cell response specificity in the premotor cortex [37] and may indicate that converging inputs to the motor cortex terminate very broadly and are not somatotopically specific. The extraction of relevant signals to drive the appropriate effectors may be delegated to downstream elements [38, 39] or could be achieved by having activity in non-relevant sites occupy a null-space which exists in parallel to the potent space during motor execution [40].

## Acknowledgments

This study was funded by the NIH (R01 NS110901-01) grant and the Israel Science Foundation (ISF-1801/18). Additional funding was received from the ELSC support for post-doctorate fellowship (AG and SI) and through the generous support of the Baruch Foundation (YP).

## Author contributions

Conceptualization, Y.P.; Investigation, A.G., A.N., M.S., and R.H.; Formal analysis, A.G and M.S.; Writing- original draft, Y.P. A.G, and S.I.

## STAR-Methods

### RESOURCE TABLE

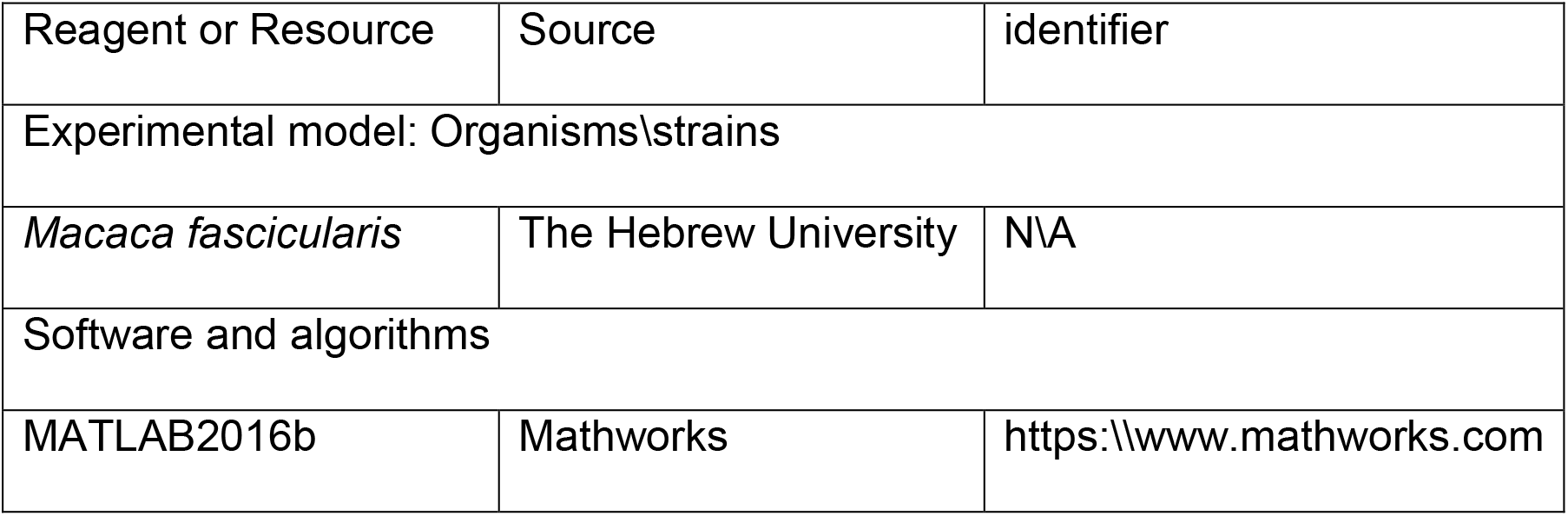

| Reagent or Resource | Source | identifier |
| --- | --- | --- |
| Experimental model: Organisms\strains |  |  |
| *Macaca fascicularis* | The Hebrew University | N/A |
| Software and algorithms |  |  |
| MATLAB2016b | Mathworks | https://www.mathworks.com |

### CONTACT FOR RESOURCE SHARING

Further information and requests for resources and reagents should be directed to and will be fulfilled by the lead contact, Yifat Prut.

### EXPERIMENTAL MODEL AND SUBJECT DETAILS

Three female monkeys (*Maccaca Fascicularis*, 3-7 Kg) were trained to perform two different 2-dimensional motor tasks: an isometric wrist task (WRIST task) and a reaching task that required shoulder-elbow movements (REACH task). Two other monkeys were trained on one of these tasks, either REACH or WRIST. All the procedures for monkey care and surgery were in strict compliance with the Hebrew University Guidelines for the Use and Care of Laboratory Animals in Research and the study was approved by the Hebrew University ethics committee.

## METHOD DETAILS

### Behavioral paradigms

#### Isometric wrist task (WRIST)

Monkeys were trained to control an on-screen cursor by applying a 2D isometric torque at the wrist (flexion/extension and radial/ulnar), while the hand of the monkey was secured by a cast in a pronation position. The task (**Fig. 1A**) included a pre-cue and delay periods in which the monkey was required to keep the cursor in the central target by applying zero torque. At the “*Go*” signal the monkey was required to move the cursor to the previously highlighted target (*Cue*) by exerting a wrist torque with the appropriate direction and amplitude and keeping the cursor in the target for the *Hold* period. Correct performance resulted in a reward (a small drop of applesauce).

#### Reaching task (REACH)

In this task (**Fig. 1B**) monkeys wore a KINARM exoskeleton (BKIN Technologies Ltd., Canada) and made planar shoulder-elbow reach movements. On each trial the monkey kept the cursor within the central target during a pre-cue and delay period and then, upon receiving the “Go” signal, had to reach a peripheral target for a reward. In this case, reaching was made via flexion/extension of shoulder and elbow joints.

### Data acquisition

After a period of training on each task, a cortical chamber was implanted above the motor cortex. After a period of recovery and retraining, extracellular recordings of motor cortical activity were made. For reach recording session, we inserted recording electrodes (1-4 individually moveable glass-coated tungsten electrodes, impedance 300-800 kΩ at 1kHz) at different cortical sites in the sensorimotor cortex. The raw signal (gain x10K, filter 300-6000 Hz) was digitized at 25-32Khz.

### Cortical mapping

Mapping was performed at the end of each recording day with the arm and the hand of the monkey freed from the cast to allow inspection of joint movements in response to the stimulation train (50 ms of biphasic stimulation delivered at 300 Hz with intensity ≤60 µA). The observed motor response was used to identify the body part controlled by the local area (shoulder, elbow, fingers, wrist, etc.). The threshold level at which an observable response was obtained, together with the anatomical landmarks were used to define the Premotor/M1 boundary. In general, sites that were close to the central sulcus and had a threshold for evoked response <10μA were referred to as M1. More rostral sites with thresholds higher than 10μA were defined as premotor sites.

### Time sequence of training and recording

Three monkeys (T, Y and M) were trained on the two different tasks in different orders (Fig. 1C-E). All monkeys performed the task with their right hand. Monkey T and monkey D were first trained on the REACH and then on the WRIST task, whereas monkey Y was trained in the reverse order. For all three monkeys, the transition to the new paradigm required a retraining period that was followed by a recording period. The exact times allotted to training, recording and retraining are shown in figure 1C-E.

One of the three monkeys (monkey T) had its implant re-secured in the transition between REACH to WRIST tasks and its chamber was replaced by a larger chamber, though the relations to the original chamber were measured.

Two additional monkeys were trained on one task alone. Monkey V was trained for 12 months on the WRIST task after which recording lasted for 9 months. Monkey C was trained for 8 months on the REACH task performed with its right hand after which recordings lasted for 4 months. The monkey was then retrained to work with its left arm for 5 months and recordings from the contralateral hemisphere lasted for 5 months. Since we found no differences in the representation between the two datasets obtained for this monkey, data from both hemispheres were pooled.

### Analyses of single cell response properties

We used offline sorting techniques (Alphasort by Alpha Omega, Nazerath Israel) to extract single cells from the compound signal. For each cell we computed its preferred target and the significance level of the tuning (Shalit et al. 2010). For cells that were significantly tuned we computed the Peri-Event Time Histograms (PETHs) for the preferred target ± 1 neighboring target.

